# Nuclear-cytoplasmic compartmentalization promotes robust timing of mitotic events by cyclin B1-Cdk1

**DOI:** 10.1101/2021.07.28.454130

**Authors:** Gembu Maryu, Qiong Yang

## Abstract

Studies applying well-mixed cytosolic extracts found the mitotic network centered on cyclin-dependent kinase (Cdk1) performs robust relaxation oscillations with tunable frequency [1–6]. However, recent work also highlighted the importance of cyclin B1-Cdk1 nuclear translocation in mitotic timing [7, 8]. How nuclear compartmentalization affects the oscillator properties and the accurate ordering of mitotic events, especially in embryos lacking checkpoints, remains elusive. Here we developed a Förster resonance energy transfer (FRET) biosensor for analyzing Cdk1 spatiotemporal dynamics in synthetic cells containing nuclei compared to those without. We found cellular compartmentalization significantly impacts clock behaviors. While the amplitude-frequency dependency measured in the homogeneous cytoplasm showed highly tunable frequency for a fixed amplitude, confirming predictions by non-spatial models [4], the frequency remains constant against cyclin variations when nuclei are present, suggesting a possible buffering mechanism of nuclear compartments to ensure robust timing. We also found all cyclin degrades within similar mitotic durations despite variable interphase cyclin expression. This scalable degradation of cyclin may further promote the precise mitotic duration. Simultaneous measurements revealed Cdk1 and cyclin B1 cycle rigorously out of phase, producing a wide orbit on their phase plane, essential for robust oscillations. We further mapped mitotic events on the phase-plane orbits. Unlike cytoplasmic-only cells showing delayed Cdk1 activation, nucleus-containing cells exhibit steady cyclin B1-Cdk1 nuclear accumulation until nuclear envelope breakdown (NEB) followed by an abrupt cyclin-independent activation to trigger anaphase. Thus, both biphasic activation and subcellular localization of Cdk1 ensure accurate ordering of substrates.

## Results & Discussion

### EKAREV FRET biosensor was modified for cyclin B1-Cdk1 activity monitoring

A Cdk1 FRET biosensor was previously developed based on a polo-box domain (PBD) of Plk1 as the phospho-binding domain and the autophosphorylation site from human cyclin B1 [9] and applied in mammalian cells [8–10] and *Drosophila* embryos [11]. However, it did not perform well in our system using *Xenopus* extracts, probably due to the varied sensor domain specificity in different species as the trough-to-peak FRET/CFP ratio was worse in *Xenopus* (less than 1%) (Figure S1B) and *Drosophila* (6%) [11] compared to the mammalian system (10-15%) [9]. The poor solubility of this protein expressed in *E. coli* is another challenge of using this sensor in cell-free systems [12]. A modified version by replacing PBD with the FHA2 domain was also reported in Hela cells but showed substantial cell-to-cell variability in the emission ratio [12]. On the other hand, an optimized backbone structure has been demonstrated to significantly reduce the basal FRET/CFP ratio and increase the gain of various intramolecular FRET biosensors, including the extracellular signal-regulated kinases (ERK) FRET sensor. The key components that make this structure superior to the prototype biosensors are a long, flexible EV linker and the optimized FRET donor and acceptor pairs. To design a new Cdk1 sensor with an improved signal-to-noise ratio, we adapted as a template an existing ERK FRET biosensor that has been optimized with an EV linker (EKAREV) [13, 14] and modified its substrate sequence to visualize a mitotic signal. The resulting sensor (hereafter named Cdk1-EV) includes a donor-acceptor fluorescent protein pair with optimized energy transfer efficiency (SECFP and YPet), a phosphorylation sequence from Cdc25C, a substrate of Cdk1 (sensor domain), the WW domain as the phosphopeptide binding domain (ligand domain), and flexible 116 amino acid EV linker (Figure 1A). The substrate sequence from Cdc25C serves as a consensus phosphorylation site for Cdk1 [15, 16] and is phosphorylated in the mitotic phase [17]. Upon cyclin B1-Cdk1 activation, the sensor domain is phosphorylated by Cdk1 and bound by the ligand domain, thereby inducing a conformational change that alters the FRET efficiency between fluorescent proteins that can be quantified by calculating ratio values of the donor to acceptor emission fluorescence intensities. To improve the specificity for the Cdk1 kinase activity, we removed the ERK-specific binding sequence FQFP peptide from the sensor domain of Cdc25C (Figure 1B).

**Figure 1.**
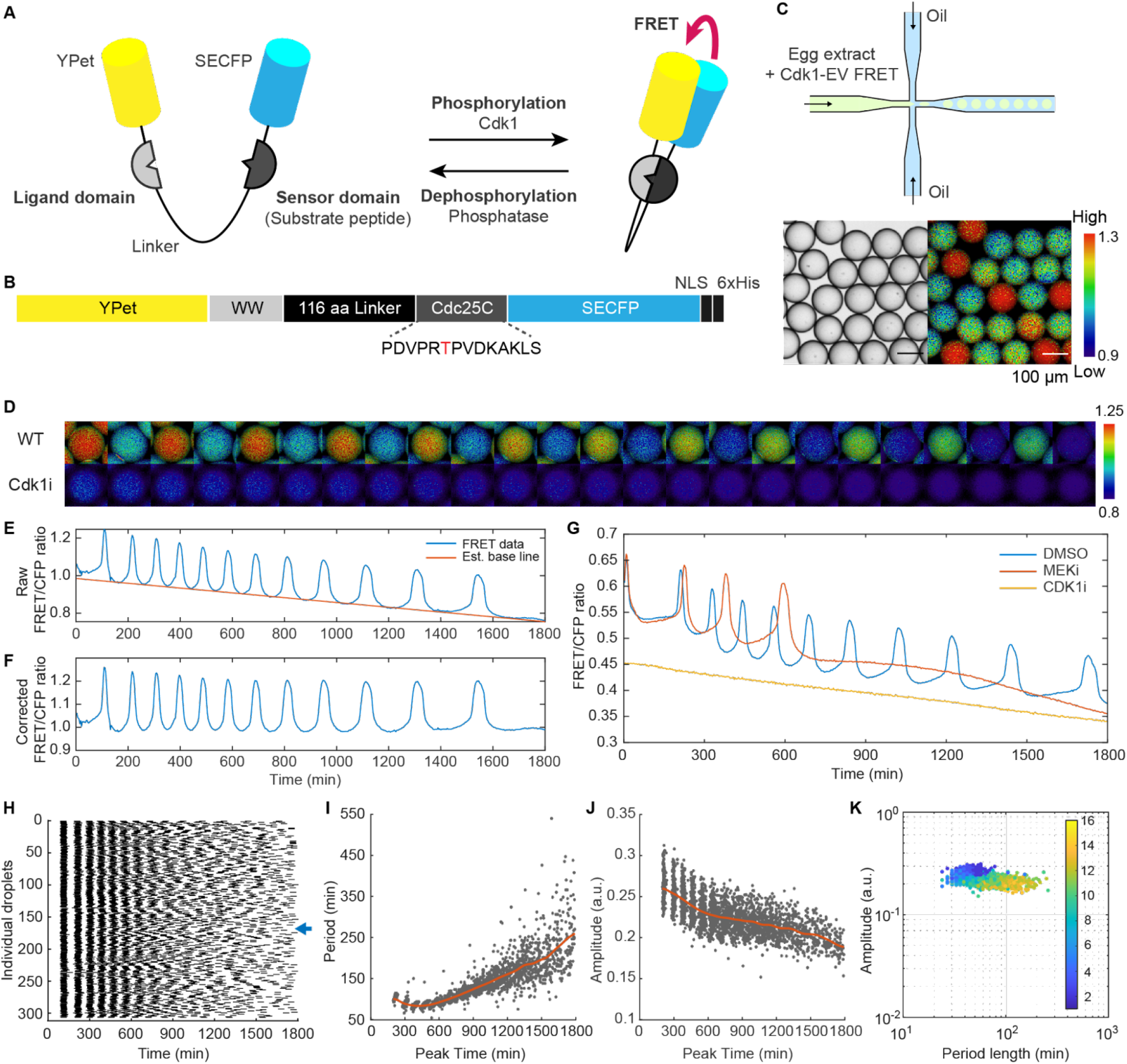
The Cdk1-EV detects oscillations of cyclin B1-Cdk1 kinase activity specifically. (A) Scheme of Cdk1-EV FRET biosensor. The FRET efficiency increases by virtue of phosphorylation of the substrate sequence of Cdk1. The phosphorylation state of the biosensor is reversible by the antagonistic phosphatase (e.g. PP2A). (B) The substrate sequence was modified from EKAREV by removing the ERK-specific binding domain. Threonine (red) is a phosphorylation site by Cdk1. The biosensor was tagged by the C-terminus Histidine tag for protein purification. (C) Top: Cycling *Xenopus* egg extracts were supplemented with recombinant Cdk1-EV FRET biosensor protein. FluoroSurfactant-stabilized droplets were formed uniformly on a microfluidic device with fluorinated oil. Bottom: A bright-field image was used for droplet segmentation in the image processing process. A representative emission ratio image was shown in the intensity-modulated display (IMD) mode. The color bar represents low (blue) to high (red) Cdk1 activity. The scale bar is 100 μm. (D) Representative droplets under positive control (WT, top row) and 1 μM Cdk1/2 inhibitor III treated condition (Cdk1i, bottom row). The emission ratio images of WT in the mitotic phase (peak) and interphase (trough) were shown alternately. Cdk1i images were selected at the same frame as WT. (E) Raw emission ratio timecourse of the WT droplet shown in (D) (blue). A Cdk1 activity baseline was estimated with trough values of Cdk1 oscillations (red). (F) Same as (E) with baseline decay corrected. Corrected data were used to quantify the amplitude of the rising and falling phases in later analysis. (G) Representative plots under inhibitor treatment conditions. 1 μM MEK inhibitor PD0325901 (red), 1 μM Cdk1 inhibitor Cdk1/2 inhibitor III (yellow), and DMSO (blue) as a vehicle of compounds were added to egg extract. FRET/CFP ratio decreased even though there was no oscillation under the Cdk1 inhibitor added condition. In MEK inhibitor treated droplets, the number of cycles was decreased and their periods were longer than the positive control condition. (H) Raster plot of FRET/CFP ratio peaks over time for over 300 individual droplets combined from multiple positions. The droplets were initially synchronized and dephased at later cycles. The blue arrow indicates the representative droplet shown in (D-F). (I, J) Period length (I) and amplitude (J) of Cdk1 oscillation over peak time. The trend was shown by kernel density estimation (KDE) (red). (K) The fold change of period length was greater than amplitude. The color bar represents the cycle number.

### The biosensor is specific to cyclin B1-Cdk1 kinase activity

To test the Cdk1-EV FRET biosensor, we purified the protein by bacterial overexpression system to apply to *Xenopus* cycling egg extract before encapsulating them in cell-sized microemulsion droplets via a microfluidic device [18–20] (Figure 1C). We observed that the FRET/CFP emission ratio changed periodically in each droplet (Figure 1D, Top row; Figure 1E, Blue; Video S1) despite an overall decay of the baseline (Figure 1E, Red). The baseline decay was likely caused by photobleaching rather than some intrinsic signal reduction, as it was also seen in the phospho-dead TA mutant (Figure S1A, Red) and under Cdk1 inhibitor-treated condition (Figure 1G, Yellow). After baseline correction, the FRET/CFP ratio reports 12 undamped, self-sustained oscillations between active and inactive states of Cdk1 (Figure 1F). The emission ratio from trough to peak of this Cdk1-EV biosensor was above 20%, better than the reported values for the Plk1-cyclin B1-based Cdk1 FRET sensor in mammalian cells (10-15%) [9] and *Drosophila* embryos (6%) [11]. We also compared the two sensors side-by-side in our cell-free system prepared from the same batch of eggs. While the Cdk1-EV sensor reported clear oscillations (Figures S1B-C, Blue), the Plk1-cyclin B1-based Cdk1 sensor could not regardless of its original linker (Figure S1B, Red) or a long, flexible EV linker (Figure S1C, Red). These extracts themselves are oscillatory as the reconstituted nucleus can undergo periodic nuclear envelope breakdown and reformation (Figures S1A-C, Right panels).

To demonstrate Cdk1-EV sensor responds only to cyclin B1-Cdk1 kinase activity, we treated the same extract with DMSO (positive control), 1 μM Cdk1/2 inhibitor III (CAS 443798-55-8), or 1 μM PD0325901 MEK inhibitor before encapsulation. In contrast to oscillations observed in the DMSO control (Figure 1G, Blue), the emission ratio in the Cdk1 inhibited droplets was completely suppressed to a constantly low level, as expected from inactive Cdk1 (Figure 1D, Bottom row; Figure 1G, Yellow). Nevertheless, MEK inhibition did not terminate oscillations (Figure 1G, Red). It is plausible that it showed a longer period than the control because ERK is one of the pivotal kinases for cell cycle progression [21]. Moreover, we did not observe spontaneous ratio change in interphase as EKAREV reported in human culture cells [17, 22]. These results indicate the Cdk1-EV sensor specifically measures the Cdk1 kinase activity with higher sensitivity than the previously reported sensor.

### Cell cycle behaves as a period-tunable, amplitude-constant oscillator in non-compartmentalized cytoplasm

With this sensor, we first examined the cell cycle amplitude-frequency dependency in these droplet cells. The frequency and amplitude are functionally essential properties of biological clocks. While it is hard for a single negative feedback loop (e.g., Goodwin oscillator) to vary its frequency without affecting its amplitude, theoretical studies suggested positive feedback can allow a broad frequency tuning for a constant amplitude, a phenomenon called frequency tunability shared by various oscillators like heartbeats and neuronal spiking [4]. Although the Cdk1 network dissection in *Xenopus* extracts has identified interlinked positive feedback of Cdk1/Wee1/Cdc25 [23, 24] and negative feedback of Cdk1 with the anaphase-promoting complex or cyclosome (APC/C) [2, 3, 5, 25], there have been no such measurements to date to confirm its frequency tuning decouples from amplitude. Bulk extracts cannot precisely recapitulate the relation as the ensemble average may blur the individual dependencies.

Our microfluidic system enables statistical single-cell amplitude-period measurements by manipulating and tracking oscillations in hundreds of cell-like droplets per position. In an unperturbed system, we observed initially synchronized oscillations among individual droplets were gradually dephasing over the cycles (Figure 1H), possibly contributed by both the partitioning effects during encapsulation [6, 26] and the random phase diffusion due to free-energy dissipation per cycle [27]. The oscillation period elongated significantly from roughly 50 to 300 min (Figure 1I), which we attributed to energy depletion [18], while the amplitude of the Cdk1 FRET/CFP emission ratio remains relatively constant, decreasing slightly from 0.25 to below 0.2 throughout the experiment spanning about 30 hours (Figure 1J). The trend is better visualized in an amplitude-period log-log plot (Figure 1K), showing a significantly greater fold change in period than in amplitude. The slight slope may be associated with changes in energy or other complicated factors over time that affect the system status globally, but not a direct amplitude-period coupling.

To confirm, we explicitly tuned the frequency by systematically perturbing the endogenous cyclin B mRNA translation using morpholinos (MOs), antisense oligonucleotides against the isoforms of *Xenopus* cyclin B1 and B2 mRNA species. We co-injected the MOs and a purified BFP as a MO concentration indicator into one inlet of a two-channel tuning microfluidic device before extract encapsulation, creating a broad spectrum of MO concentrations across the droplets (Figure 2A). These droplet cells displayed different patterns of oscillations (Figure 2B), having fewer cycles (Figure 2C), steadily increased rising periods (Figure 2D, Blue), and stable falling periods (Figure 2D, Red) for increased MO concentration. However, both rising and falling amplitudes were almost uniform across a broad range of MO concentrations (Figure 2E). The amplitude-period log-log plot showed a zero-slope relation, with MO-independent, narrowly-distributed amplitude versus a widely distributed period that depends on MO concentration (color-coded by the BFPintensity) (Figure 2F), indicating the decoupled frequency tuning from the amplitude.

**Figure 2.**
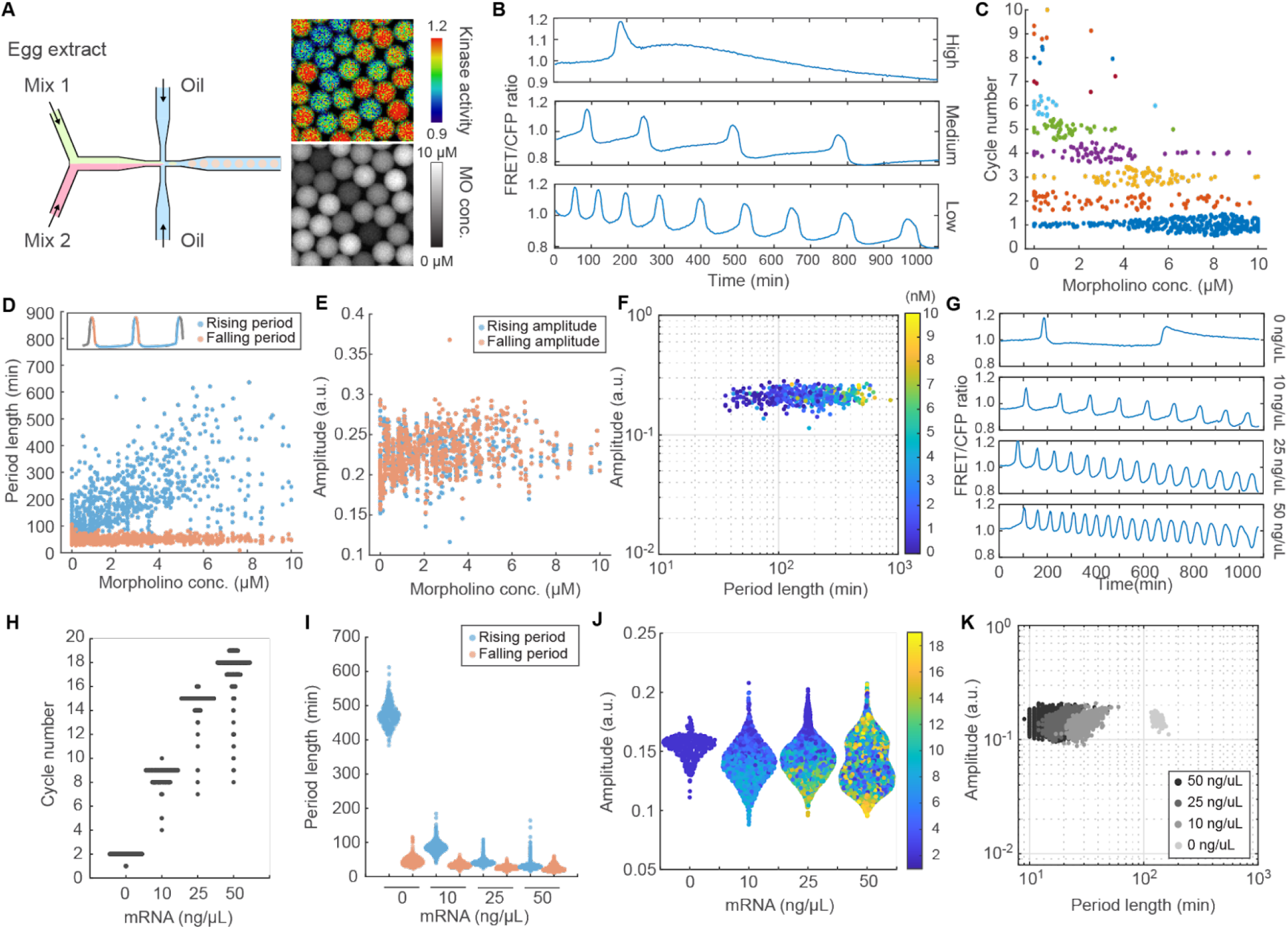
A period-tunable, amplitude-robust oscillator property was revealed by Cdk1-EV FRET measurement. (A) Scheme of two-channel tuning. Mix 1 (Extracts plus Cdk1-EV biosensor) and Mix 2 (Mix 1 plus 10 μM MOs plus recombinant BFP as a dye indicator for MO concentration) were mixed in a microfluidic device. The emission ratio for Cdk1 activation and BFP intensity of representative droplets were shown in IMD mode and gray-scale images. (B) Representative Cdk1 oscillation raw-data profiles for different morpholino concentrations. (C) The distribution of the total cycle number of individual droplets was changed with MOs concentration. The maximum total cycle number decreased with MOs concentration. Each dot indicates a single droplet. MOs concentration was estimated from BFP intensity. (D) Rising period (the time from a trough to the next peak, blue) and falling period (the time from a peak to the next trough, red) of Cdk1 oscillation versus MOs concentration. (E) Rising amplitudes (blue) and falling amplitudes (red) versus MOs concentration. (F) Amplitude (calculated as the average of rising amplitude and falling amplitude) remains constant with a wide range of period lengths. The color bar represents MOs concentration. (G) Raw Cdk1 oscillation timecourses of representative droplets containing extracts treated with 10 μM *Xenopus* cyclin B MOs and various amounts of manually added exogenous human cyclin B1 mRNA. Suppressed Cdk1 oscillation by MOs was recovered by human cyclin B1 mRNA. (H) Cycle number on average is increased with the concentration of mRNA. Each dot represents the total cycle number of a single droplet. (I) The falling period is constant across different exogenous mRNA concentrations. The rising period length is dramatically shortened as the mRNA concentration increases. (J) The range of amplitude was kept at a similar level across different mRNA concentrations. The color bar represents the cycle number. (K) Amplitude-period log-log plot for clusters of droplets with different mRNA concentrations.

We also tuned up the speed by adding human cyclin B1 mRNAs (0, 10, 25, 50 ng/μL) to the MO-treated extracts with endogenous *Xenopus* cyclin B silenced and found the Cdk1 activity waveform is closely dependent on the exogenous mRNA concentration (Figure 2G). With increased human cyclin B1 mRNA concentration, we observed cycle numbers increased (Figure 2H), rising periods decreased (Figure 2I, Blue), and falling periods invariant (Figure 2I, Orange). The amplitude was kept within a similar range across different mRNA concentrations (Figure 2J), resulting in a flat amplitude-period dependency (Figure 2K). Interestingly, tuning the system outside of the physiological range by excessive amounts of cyclin B MO (Figure S2A, Red) or mRNA (Figure S2A, Blue) will arrest the system at a stable steady-state of inactive Cdk1 (emission ratio below 1) or active Cdk1 (emission ratio above 1.2), respectively.

Together, these measurements revealed consistent clock behaviors across all different experimental conditions, e.g., highly tunable interphase duration, robust mitotic phase duration, and constant amplitude of Cdk1 activity. These are intrinsic properties associated with the Cdk1 coupled positive and negative feedback topology, as predicted by the mathematical models [4]. Our amplitude-period measurements in the cytoplasm-only cells provide the first explicit experimental evidence for this long-speculated theory.

### Nuclear-cytoplasmic compartmentalization promotes robust mitotic timing

However, recent studies have identified spatial positive feedback that drives abrupt cyclin B1-Cdk1 nuclear translocation upon activation [7], raising the question of whether the oscillator properties revealed in the spatially homogeneous extracts still hold in cells containing nuclei. To understand the impact of cellular compartmentalization, we supplied cycling extracts with demembranated sperm chromatin before encapsulation to induce the self-assembly of nuclei in droplets. Due to random encapsulation following Poisson statistics, droplets prepared from the same bulk extracts may or may not contain a nucleus, providing a unique system to compare side-by-side the clock dynamics when nuclei are absent or present (Figure 3A; Video S2). We detect nuclei in droplets using a nuclear maker, protein of nuclear localization signal (NLS) fused with mCherry (Figure 3A, Rows 2 and 4). Since activated cyclin B1-Cdk1 immediately translocates into the nucleus rapidly and irreversibly until nuclear envelope breakdown (NEB) [7, 8], we also tagged the FRET biosensor by NLS to visualize nuclear-specific Cdk1 activation efficiently. In both droplets with and without a nucleus, we observed Cdk1 kinase activity oscillations with period elongation over the cycles (Figure 3A, Row 1 and 3; Figure 3B). However, the nucleus-containing droplet showed a shorter period in each cycle than the cytoplasm-only droplet, despite that they came from the same bulk extracts. This difference is statistically significant (Figure 3C). One possible explanation for this observation may be the nucleus concentrating effect since the abrupt translocation of cyclin B1-Cdk1 from the cytoplasm into the nucleus can effectively make it more concentrated, which in turn promotes Cdk1’s nuclear substrate phosphorylation and accelerates mitotic entry. However, this would not be as effective if cyclin B is excessive in the system.

**Figure 3.**
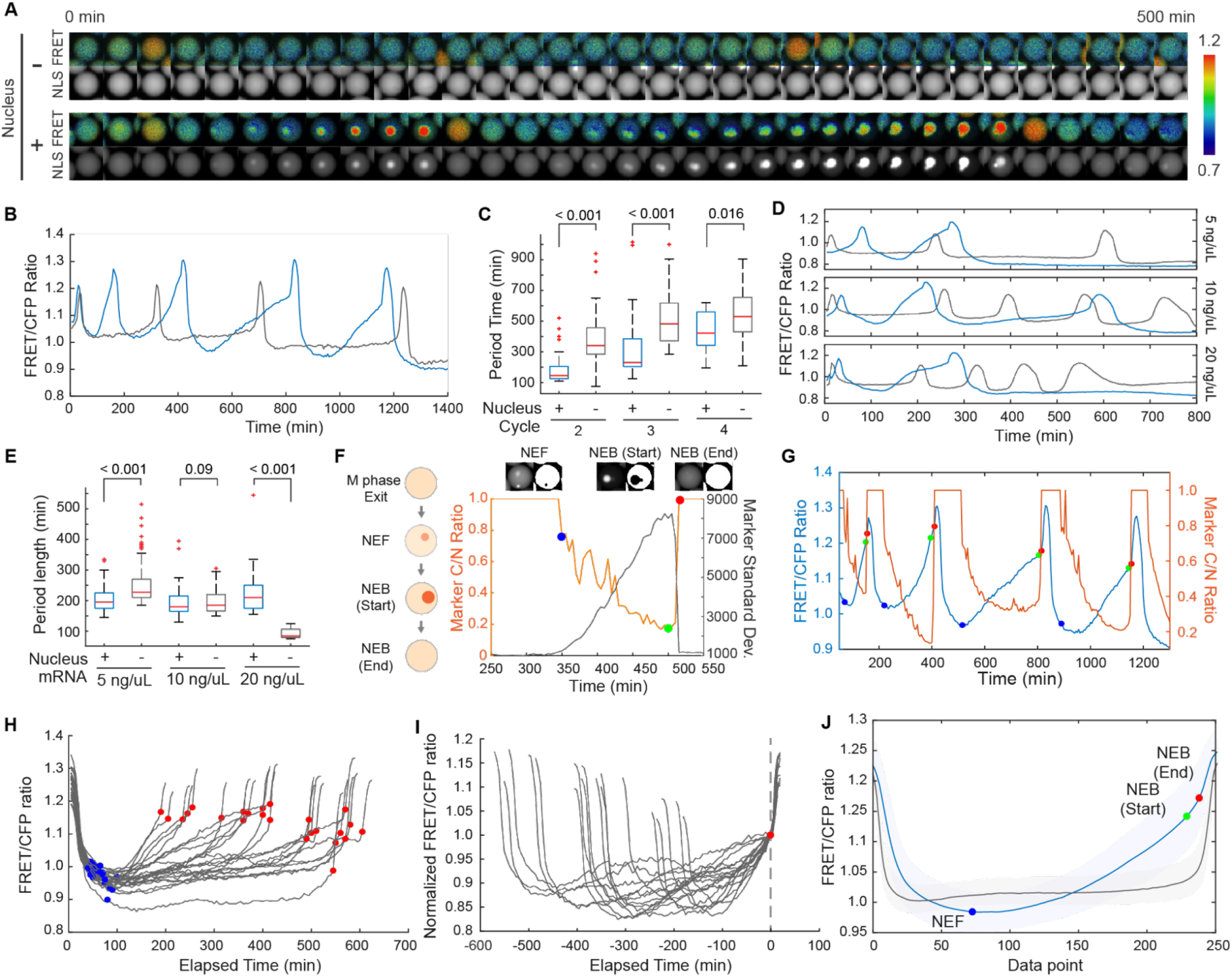
Nuclear compartmentalization affects the dynamics of Cdk1 activation and cell cycle period. (A) A representative droplet having a nucleus (Bottom panel) or not (Top panel). Cdk1 activity was represented by FRET/CFP ratio IMD images. A nucleus was detected by recombinant NLS-mCherry-NLS proteins as a bright spot in the nucleus-containing droplet. (B) Cdk1 activity timecourses for droplets in (A), showing a single spiky Cdk1 activation for the droplet with no nucleus (gray) as compared to a gradual Cdk1 activation followed by a spike for the droplet with a nucleus (blue). Ratio value was normalized by division with the average of 1st trough values of cytoplasm-only cells. (C) Period length of nucleus-containing droplets was shorter than that of cytoplasm-only droplets in each cycle. The period is calculated based on the peak-to-peak time (Cycle 2: 1st and 2nd peak, Cycle 3: 2nd and 3rd peak, Cycle 4: 3rd peak and 4th peak). Later cycles had longer period durations. (D) Representative raw Cdk1 activity of nucleus droplets (blue) and cytoplasm-only droplets (gray) with additional human Cyclin B1 mRNA. Periods of cytoplasm-only droplets were changed by additional mRNA concentration, but that of nucleus droplets was in a similar range. Also, a small jump just before the peak was observed in nucleus droplets as we showed in (B). (E) Comparison of period length difference between nucleus- and non-nucleus-droplets under different additional mRNA concentrations. Only the first oscillation was calculated. With 5 ng/μL mRNA added, nucleus droplets had shorter cycles than non-nucleus droplets. However, with 20 ng/μL additional mRNA, it showed an opposite feature. Statistical significance was not observed with 10 ng/μL mRNA. (F) The nuclear region inside the droplet was segmented based on the standard deviation value of the nuclear marker (gray) and the skewness of the nucleus marker intensity histogram (Figure S3A). The segmentation results provided us the cytosolic to nuclear (C/N) ratio value of the nuclear marker (orange). When there is no nucleus (after NEB and before NEF), C/N turns to be 1.0. Based on this value, the time point of NEF (blue), NEB Start (green), and NEB End (red) were defined as shown in representative images (Top panel, nuclear marker: left, segmentation result: right). (G) Defined nuclear envelope-related events (NEF: blue dot, NEB Start: green dot, NEB End: red dot) were plotted on Cdk1 activity (blue line) and nuclear maker C/N ratio value (orange line). Short steeper Cdk1 activation started at the NEB End. Ratio value was normalized by division with the average 1st trough value of cytoplasm-only cells. (H) Cdk1 activity aligned at the left peak (gray) was plotted with NEF (blue dot) and NEB End (red dot) information. NEF was observed with similar ratio values around 40-100 min after the aligned peak regardless of the period length. NEB End also observed similar ratio values among different period lengths. Ratio value was normalized by division with the average 1st trough value of cytoplasm-only cells. (I) Oscillation data shown in (H) were aligned at NEB End (red dot, vertical gray line) and their values were normalized by the activity at NEB End. The steepness of the ratio change after NEB was similar among droplets. (J) Cdk1 activities of both conditions were transformed into 250 datapoint arrays and their average values and their standard deviation values were plotted as solid line and shade. The time information of NEF (blue dot), NEB Start (green dot), and NEB End (red dot) was also transformed and plotted with nucleus data (blue). After NEB End, transformed data also changed its steepness. On the other hand, without a nucleus, droplets remained flat in Cdk1 activity in interphase before abrupt activation just before its peak (gray). In terms of the inactivation of Cdk1 activity, nucleus droplets were more deeply inactivated after mitotic exit than cytoplasm-only droplets. Ratio value was normalized by division with the average 1st trough value of cytoplasm-only cells.

As a test, we supplied exogenous human cyclin B1 mRNAs in addition to the endogenous mRNAs in extracts and found the difference in the periods of nuclear-containing droplets and their cytoplasmic-counterparts is reduced as the exogenous cyclin B1 mRNA concentration increases (from 5 ng/μL to 10 ng/μL) until the order switched (20 ng/μL) (Figures 3D and 3E). Intriguingly, while cyclin B can easily tune the frequency of the Cdk1 oscillator in a homogeneous cytoplasmic environment (Figure 3E, Columns 2, 4, 6), a frequency-tunability phenomenon as observed before (Figure 2I), we found the system hardly reacts to the cyclin B variation and maintains an almost constant period when a nucleus is present (Figure 3E, Columns 1, 3, 5). This robust timing could be explained by the nuclear accumulation of cyclin B1-Cdk1 against cytoplasmic fluctuations, such that when cyclin B is dilute in the cytoplasm, nuclear concentrating promotes mitotic entry but excessive cyclin B cannot further accelerate mitotic entry if the nucleus imports a fixed amount of active cyclin B1-Cdk1 before NEB. The cytoplasm in living cells is a highly fluctuating environment that can significantly reduce its density from mitotic swelling [28, 29]. Our results suggest the nucleus may provide spatial insulation from the noisy cytoplasm by compartmentalizing activated cyclin B1-Cdk1 from the rest of unbound and bound Cdk1 molecular species.

### Nuclear-cytoplasmic compartmentalization facilitates biphasic activation of Cdk1 and temporal ordering of mitotic events

We also found that the oscillation profiles are qualitatively different between droplets with or without a nucleus. Cdk1 in the cytoplasm-only droplet remains inactivated by Wee1 throughout the interphase before a sharp activation at mitosis. This delayed, spiky-shaped Cdk1 activation is well-described in cell-cycle models considering no spatial variability [5, 30] (Figure 3B, Gray). However, in the nucleus-containing droplet, Cdk1 steadily increased since the beginning of interphase to a moderate activity before a quick jump to a distinctively higher activity (Figure 3B, Blue).

To examine whether the biphasic activation of Cdk1 may correlate with any of the downstream mitotic events, we investigated the temporal order of Cdk1 activation dynamics, nuclear envelope formation (NEF), and nuclear envelope breakdown (NEB). We developed an algorithm to automatically segment the nucleus and analyze NLS-mCherry-NLS data of each droplet (Start Methods; Figure S3B, 3C). From the time courses of mCherry cytoplasmic/nuclear (C/N) ratio (Figure 3F, Orange) and standard deviation (Figure 3F, Gray), we extracted the time of NEF (Blue dot), NEB Start (Green dot), and NEB End (Red dot). We found NEF started before Cdk1 was fully inactivated, and the transition point of the two-phase Cdk1 activation coincided with the end of NEB (Figure 3G). The interphase Cdk1 activation was roughly linear with time, rate-limited by cyclin B1 production. Aligning all the individual cycles based on peak time, we found that the steepness of the first-phase Cdk1 activation is highly variable, likely caused by the cyclin B mRNA variation (Figure 3H). Yet, the activation slope is anti-correlated with the interphase period length, suggesting a robust threshold of Cdk1 activity (Figure S3D, S3E), beyond which may trigger a mitotic event, which we found is the NEB End (Figure 3H, Red dot).

Interestingly, despite the large variability of interphase activation speed, the steepness in Cdk1 second-phase activation after NEB was uniform across all droplets and collapsed into one line after aligning all time-courses based on NEB End (Figure 3I). The second activation is for only a brief period independent of cyclin B1 (10-15 min regardless of the interphase length), likely controlled by the firing of the positive-plus-negative feedback loops (analogous to the abrupt discharging of a capacitor). It may also be contributed by active cytoplasmic Cdk1 [8] accessible to NLS-tagged FRET sensors upon NEB. Regardless of which mechanism, it ensures the temporal segregation of early mitotic events NEB and late mitotic events, such as APC/C activation.

To summarize the main features of Cdk1 activation affected by compartmentalization, we standardized all data into 250 datapoint arrays (Star Methods). Without a nucleus, Cdk1 remains a constant basal level throughput interphase before a burst-like activation (Figure 3J, Gray). With a nucleus, after a deeper deactivation at NEF, Cdk1 undergoes gradual activation and a steeper activation separated by NEB (Figure 3J, blue). This two-phase activation of Cdk1 has not been reported by the Plk1-cyclinB1 based Cdk1 FRET sensor, possibly because of the relatively low sensitivity or blurred signal from shuffling of FRET sensor between the nucleus and cytoplasm, or system-specific difference. However, other studies also suggested that different levels of Cdk1 activity regulate orderly cell-cycle progression [31], and the level of Cdk1 activity required for mitotic entry is not sufficient for later mitotic progression [32, 33]. A study on fly embryos also found that while chromosome condensation can occur in the absence of a detectable Cdk1 activity, the prophase exit (NEB End) is a decision point and coincides with Cdk1 activation [34].

Subcellular localization of Cdk can contribute to the correct ordering of phosphorylation of different mitotic substrates [35–37]. Our study also highlights the role of compartmentalized Cdk1 in accurately ordering the timing of different mitotic events. Active Cdk1 initially accumulates in the nucleus to trigger NEB; after NEB, Cdk1 switches to a higher level to trigger APC activation. Thus, both Cdk1 subcellular localization (via compartmentalization) and the activity regulation are critical factors for the correct order of downstream events. These findings highlight the importance of cellular compartmentalization and the necessity of cross-checking studies using well-mixed extracts and cells, where the same molecular network may behave differently.

### High-resolution mapping of mitotic events on phase plane trajectories

The activation of APC/C by cyclin B1-Cdk1 leads to cyclin B1 destruction and Cdk1 inactivation, completing an essential negative feedback loop. To sustain robust oscillations, the cyclin synthesis-degradation cycle and Cdk1 activation-deactivation cycle must go out of phase to form a wide loop in the phase plane of active Cdk1 and cyclin B1 abundance. The shape of the phase-plane orbit is crucial to understand the dynamic features of the system. Theoretical studies have predicted a wide triangular-shaped loop [4, 38]. Although it has been tested experimentally in bulk extracts, the measurement was only for one cycle, and the shape could only reflect the average of orbits from an ensemble of oscillators [38]. Here, we performed a real-time tracking of the phase-plane trajectories at the single-cell level for multiple cycles, to obtain a detailed picture of the Cdk1 cycle in relationship to the cyclin cycle as well as their relations with the nuclear events. We supplied both the human cyclinB1-mScarlet-i mRNA and Cdk1-EV biosensor in droplet cells with nuclei and simultaneously measured the spatiotemporal dynamics of both cyclin B accumulation and Cdk1 activity over time (Figure 4A; Video S3). We observed colocalization of Cdk1 FRET and cyclin B1-mScarlet signals inside the nucleus until NEB, consistent with the observations of nuclear localization of cyclin B1-Cdk1 complex upon activation in the late G2 phase in mammalian cells [7, 8]. But in mammalian cells, cyclin B was localized in the cytoplasm until late G2, while we do not observe any enriched cytoplasmic localization of cyclin B1 since the rapid early embryonic cycles skip the G1 and G2 gap phases.

**Figure 4.**
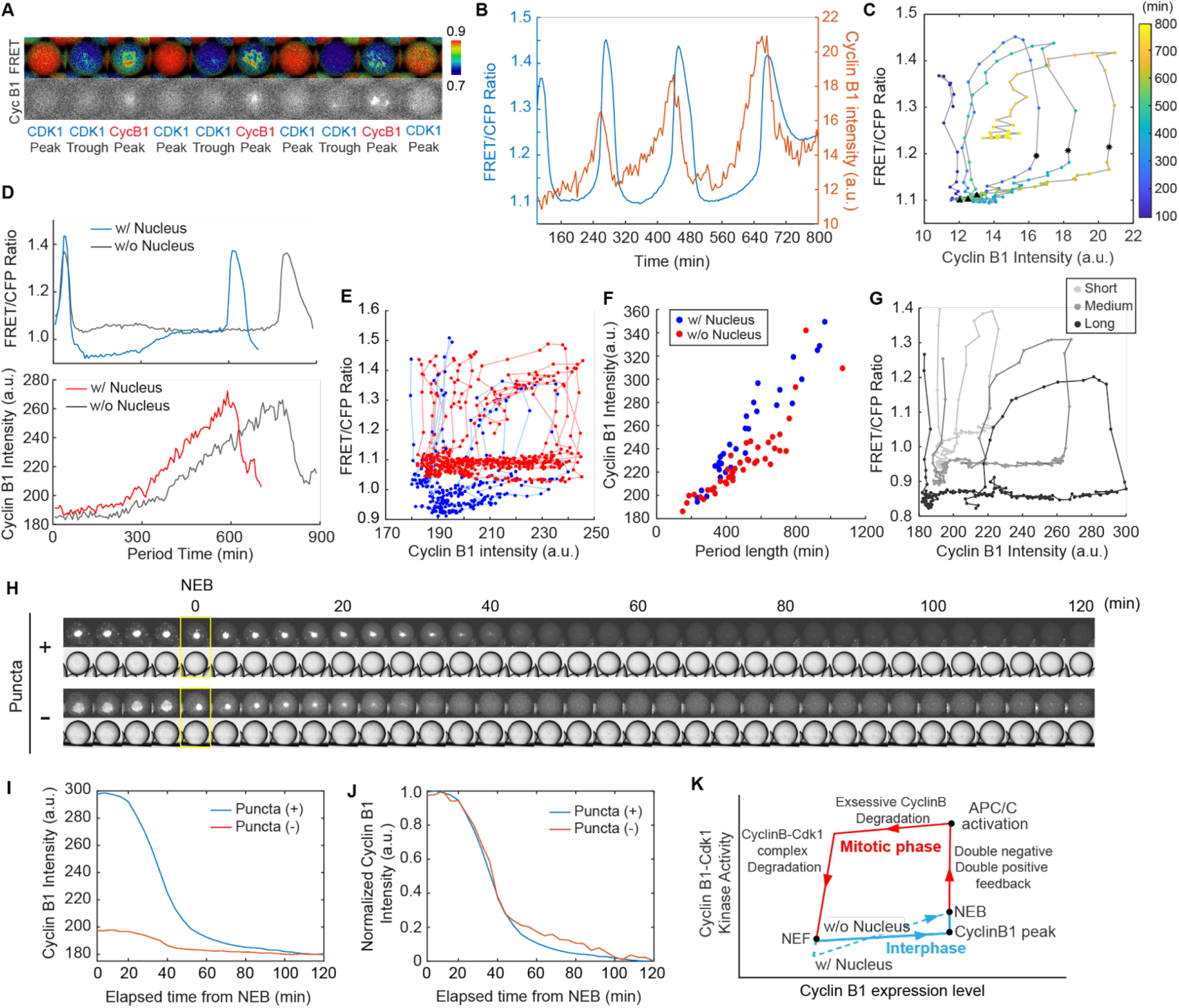
Simultaneous measurements of cyclin B1 expression and the Cdk1 activity. (A) A representative droplet was shown with FRET/CFP ratio IMD images and cyclin B1-mScarlet-i expression images. Images for Cdk1 peak, Cdk1 trough, and cyclin B1 peak were selected. There was a time gap between the peaks of Cdk1 activity and cyclin B1 expression level. This droplet was treated with 10 μM MOs and 50 ng/μL mRNA. Bright spot in the cyclin B1 channel colocalizes with that in the NLS-tagged Cdk1-EV FRET channel. (B) Both Cdk1 kinase activity and cyclin B1-mScarlet-i expression were quantified simultaneously for multiple cycles. The linear decay of FRET ratio value was corrected with linear baseline estimation shown in Figure1, and the corrected signal value was divided by the average of 1st trough values of cytoplasm-only droplets. (C) Quantified data in 4B was plotted as a phase-plane trajectory with NEF (black triangle) and NEB End (asterisk) marked. Colorbar indicates elapsed time from experiment start. Although peak and trough values of Cdk1 activity were similar among cycles, the phase-plane orbits shift to the right with time. (D) Representative droplets that had a similar cyclin B1-Katushka expression level (with 10 ng/μL mRNA added). Emission ratio amplitude was also similar between spatially different environments. The period length of the nucleus droplet (blue in upper panel, red in lower panel) was shorter than the nucleus-free droplet (gray). The linear decay of FRET ratio value was corrected with linear baseline estimation shown in Figure1, and the corrected signal value was divided by the average of 1st trough values of cytoplasm-only droplets. (E) The shapes of phase-plane orbits were compared between spatially homogeneous (red) and heterogeneous (blue) cellular environments. It was lower than that of nucleus-absent droplets (red) as described in Figure 3J. The path of Cdk1 activation after NEB and cyclin B1-Cdk1 inactivation by APC/C was not different between conditions. The linear decay of FRET ratio value was corrected with linear baseline estimation shown in Figure1, and the corrected signal value was divided by the average of 1st trough values of cytoplasm-only droplets. (F) Period length and maximum intensity of cyclin B1 expression were linearly correlated. When droplets had similar cyclin B1 expression levels under different nucleus conditions, nucleus droplets showed shorter periods as we observed in Figure 3C. (G) Representative phase-plane orbits of different period lengths were plotted for droplets with a nucleus. Phase plane orbits shifted right and became wider as period length increased. A plateau high Cdk1 activity was observed from the Cdk1 activity peak until the onset of Cdk1 inactivation in a longer period droplet. The linear decay of FRET ratio value was corrected with linear baseline estimation shown in Figure1, and the corrected signal value was divided by the average of 1st trough values of cytoplasm-only droplets. (H) Representative droplets that expressed cyclin B1-mScarlet-i protein with or without puncta in the cytosolic region under 10 ng/μL added condition. Puncta were observed in a higher cyclin B1 expressed droplet. The required time for complete degradation of cyclin B1 was comparable between different conditions. Contraction of cyclin B1 was observed after NEB (yellow square). The Time interval of the montage was 4 min. (I) Degradation trajectory of puncta droplets (blue) and without puncta droplets (red). Both droplets showed a similar background intensity after the complete degradation of cyclin B1. (J) Normalized cyclin B1 degradation trend of puncta droplets (blue) and without puncta droplets (red). Both types of droplets had a similar trend. (K) Summary of Cdk1 activity and cyclin B1 expression dynamics that were observed in this study.

The peaks and troughs of cyclin B1 expression are clearly out of phase with Cdk1 activity (Figure 4B), creating a wide loop for each cycle in the phase-plane (Figure 4C). Cyclin B1 peaks at the end of the first activation of Cdk1, but the degradation of cyclin B1 is not detectable until Cdk1 jumps to a peak, suggesting the second activation of Cdk1 is required for activation of APC/C. The phase-plane trajectory features square-shaped orbits, each containing four branches in temporal order: a slow phase of Cdk1 first activation, which is rated limited by cyclin B1 synthesis (Branch #1: bottom horizontal); an abrupt activation of Cdk1 during which cyclin B has no change (Branch #2: right verticle); degradation of cyclin B during which Cdk1 changes a little (Branch #3: top horizontal); a fast Cdk1 inactivation (Branch #4: left vertical). The progression speed can be estimated by how long it is required to pass each phase (by the number of data points per branch for 4 min each), suggesting interphase (Branch #1) is the slowest and mitotic phases are fast (Branches #2-4). We also mapped NEF (black triangles) and NEB (black asterisks) on the orbits to obtain the accurate temporal correlation of key mitotic events with cyclin B1-Cdk1 dynamics. NEB occurred immediately after cyclin B1 reached its peak and before Cdk1 second activation jumps. NEF happens when Cdk1 activity drops below a threshold that is lower than the threshold for NEB. Interestingly, Cdk1 inactivation continues even after NEF and cyclin re-accumulation, an observation revealed by high-resolution mapping.

Cyclin B1 is a rate-limiting factor for interphase Cdk1 activation, meaning that a slower cyclin synthesis usually gives a longer interphase period. However, we observed the period over the cycles elongated even though the cyclin B1 synthesis rate did not decrease, resulting in increased cyclin B peak expression (Figure 4B, Red) and right-shifted phase-plane orbit each cycle (Figure 4C). We attributed this period elongation to a harder-to-activate Cdk1 by cyclin B1 that requires a significantly larger threshold of total cyclin B in each cycle to reach the same Cdk1 activity threshold for triggering NEB (Figure 4C, Lower branches, and black asterisks). We previously predicted the right-shifted orbits and the increased threshold of Cdk1 activation by cyclin B1 via an energy consumption model [18], where the decreased ATP over time favors more unphosphorylated Wee1 (inhibitor of Cdk1) and less phosphorylated Cdc25 (activator of Cdk1), causing less Cdk1 activated for the same cyclin B1 accumulation. Our ability to trace multiple cycles of single-cell phase-plane trajectories in real-time provides an essential experimental verification for the energy-depletion theory [18].

We further explored the impact of compartments on trajectory shapes. The nucleus-present cells showed advanced mitotic entry than the nucleus-absent cells that have comparable cyclin B expression (Figure 4D, both with MO + 10 ng/μL human cyclin B1 mRNA), consistent with Figure 3B. Both showed square-shaped orbits, except for several different features in their lower branches (Figure 4E). In the nucleus-containing droplets, we observed an overshoot of Cdk1 inactivation even after cyclin B1 hits the trough, resulting in a lower basal Cdk1 activity than the nucleus-absent droplet. Additionally, interphase Cdk1 stays at a low constant in the cytoplasm-only droplets but gradually increases with cyclin B in droplets containing nuclei, resulting in flat (Red) versus oblique (Blue) branch #1. While droplets with nuclei gave shorter periods than those without for a comparable cyclin B1 expression (Figure 4F, Blue left-shifted to Red), both showed increasing cyclin peak intensities with period lengths, further supporting the idea that the rate-limiting factor cyclin B alone cannot determine the oscillation speed and is subject to the Cdk1 activation threshold, which itself can vary by energy availability.

We also compared phase-plane orbits of different period lengths, displaying a wider loop and larger cyclin B1 peak-value for a longer period (Figure 4G; Figure S4A for all oscillations). Droplets with short periods have no obvious branch #3, and Cdk1 inactivates almost immediately following the degradation of cyclin B1, exhibiting a roughly triangular-shaped orbit, supporting the previous model predictions [4, 38]. But the longer-period droplet not only produces more cyclin B (a wider branch #1) but also has more to degrade before Cdk1 starts to inactivate (a wider branch #3), suggesting that the excessive unbound cyclin B1 was degraded before the bound cyclin B1. Thus, whether the orbit is triangular- or square-shaped depends on the amount of excessive unbound cyclin B1. Consistent with our findings, studies in mammalian cells also suggested that excessive cyclin B1 must be degraded by the time of anaphase onset because even a low level of nondegradable cyclin B1 is sufficient to block the metaphase-anaphase transition [39]. Meiosis of mouse oocytes also preferentially degrades free cyclin B1 [10].

To further understand the spatiotemporal dynamics of unbound and bound cyclin, we compared droplets with distinguished cyclin B1 expressions. In droplets of over-expressed cyclin B1, we observed puncta-like structures that were not seen in the Cdk1 FRET channel (Figure 4H; Figure S4B, Video S4), suggesting these are unbound cyclin B. Similar punctate structures of cyclin B1 were also observed in the cytoplasm of starfish oocytes that disappear upon activation of Cdk1 [40]. Other than the puncta distributed in the cytosolic region, we also observed a large bright structure localized right beside the nucleus, but not inside (Figure S4C). The small puncta remained in the cytoplasm until NEB when they rapidly aggregated towards the larger structure before being degraded. Further experiments are required to understand this rapid aggregation. However, cyclin B1 is reported to localize to centrosomes, microtubules, chromatin, and kinetochores [41, 42], and the sudden contraction of cyclin B1 punta upon NEB may be achieved via active microtubule transportation for localization to kinetochores or chromatin DNA.

Despite the significantly different cyclin B1 expression in these droplets, it took a similar time for them to be completely degraded (Figure 4I). Cyclin B1 degradation is faster in the droplet with over-expressed cyclin B1. But the relative degradation trend is similar (Figure 4J), with a comparable half maximum time (the time taken to reach 50% of the peak), implying that the degradation process is in a scalable manner regardless of the absolute amount of expressed proteins. Studies in different organisms have found that mitosis can insulate itself from the highly variable interphase and remain constant, thanks to the positive feedback regulation [20, 43]. Our finding of scalable degradation also provides an essential mechanism to ensure the droplets that have high variability in cyclin B1 expression and interphase durations still maintain robust mitosis.

As illustrated in Figure 4K, our study provided a detailed mapping of phase-plane trajectories with a unique comparison between the presence and absence of a nucleus. In both cases, cell cycle functions as a typical relaxation oscillator, characteristic of a slow buildup (gradual cyclin synthesis, Blue) and a fast discharge (switching of Cdk1 in the robust mitotic phase, Red), analogous to a capacitor that is charged slowly (stress built-up) but discharged rapidly (stress relaxed). The nuclear compartment significantly impacts the Cdk1 response to cyclin during interphase, changing it from flat to an oblique increase (Blue). The cyclin-independent overshoots of Cdk1 inactivation beyond NEF and activation beyond NEB may feature hysteresis and ensure irreversible decision-making at mitotic entry and mitotic exit. These overshoots and compartmentalization of Cdk1 together govern the temporal order of its downstream mitotic events.

## Acknowledgments

We thank Michiyuki Matsuda, Dorus Gadellaand, Xuedong Liu for providing pPBbsr2-3594nls, pmScarlet-i_C1, and pXL-6xHis-CDK1-FRET-JP (Originally developed by John Pines group) constructs respectively; Yohei Kondo for providing custom ImageJ macro; Meng Sun, Shiyuan Wang, Owen Plus, and Minjun Jin for helping on the preparation of microfluidic device and *Xenopus* egg extracts. This work was supported by grants from the NSF (MCB #1817909; Early Career #1553031), NIH (NIGMS#R35GM119688), and Alfred P. Sloan Foundation.

## Author Contributions

G.M. and Q.Y. conceived the project; G.M. performed experiments and analyzed the data; G.M. and Q.Y. wrote the manuscript.

## Star Methods

**KYE RESOURCES TABLE.**
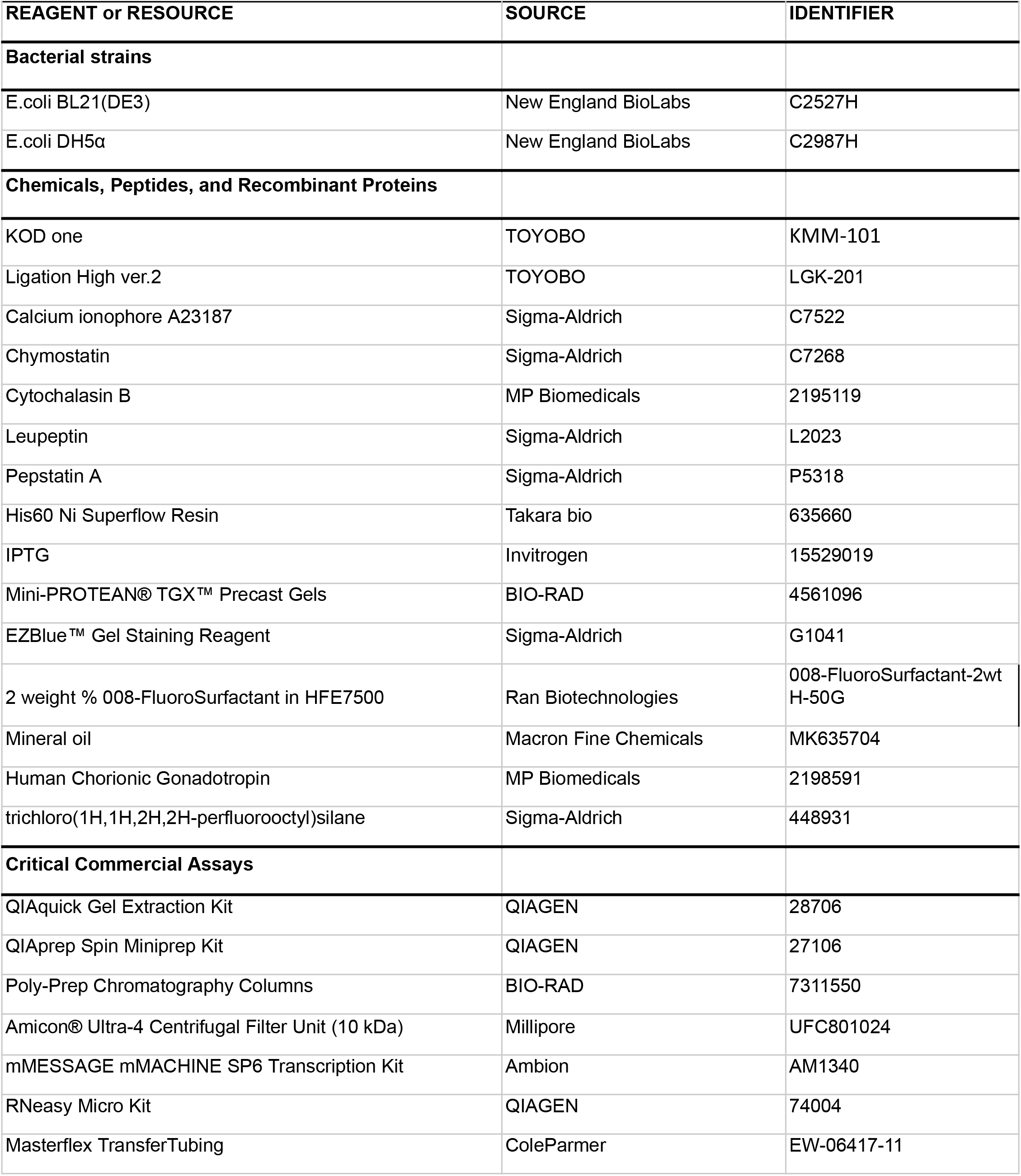
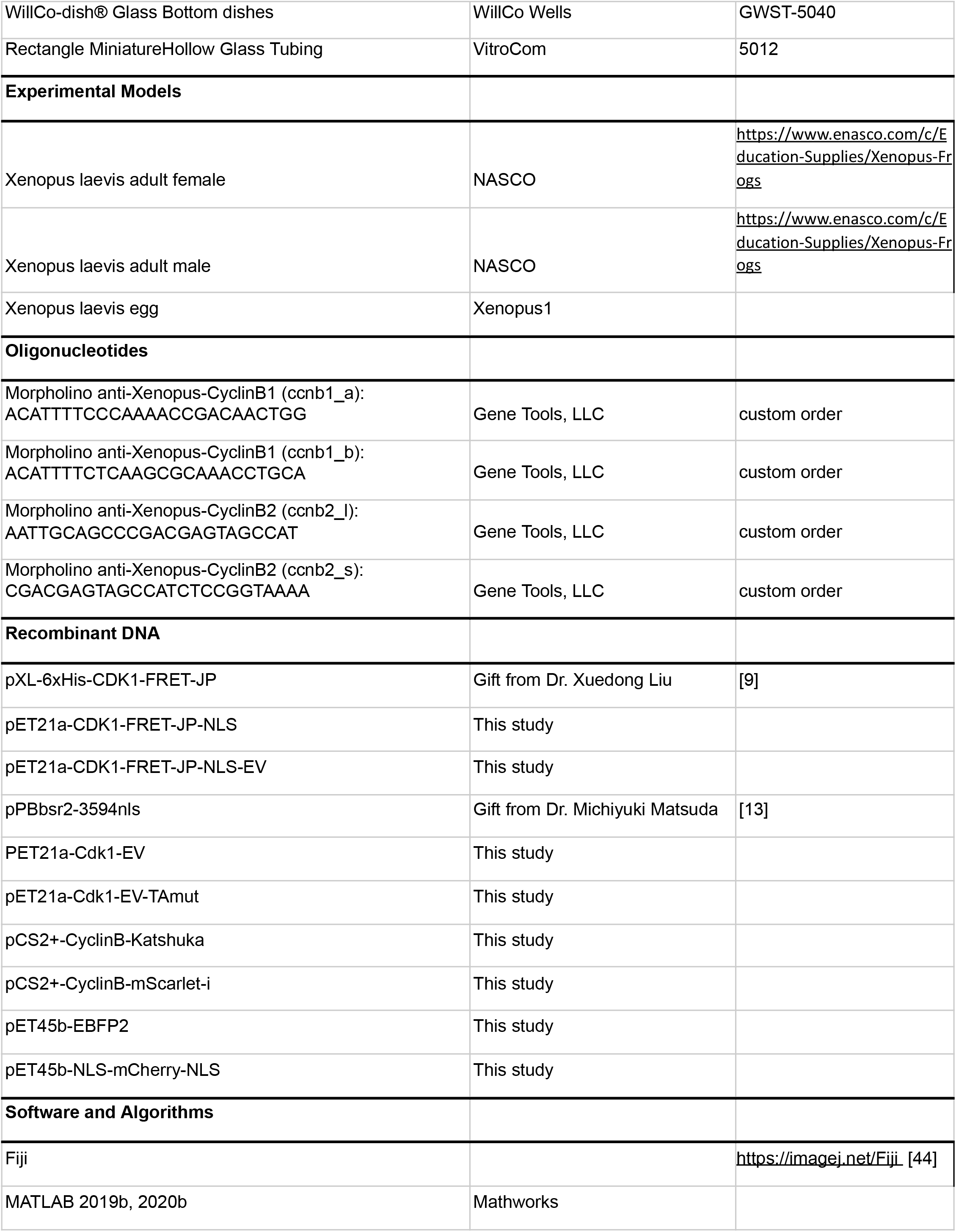
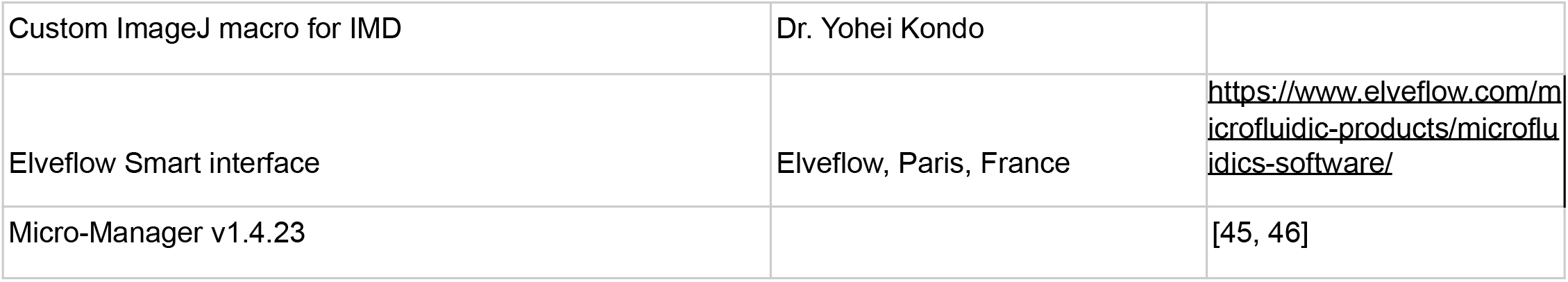

### RESOURCE AVAILABILITY

#### Lead Contact

Further information and requests for resources and reagents should be directed to and will be fulfilled by the lead contact, Qiong Yang.

#### Material Availability

Plasmids generated in this study can be obtained through the Lead Contact.

#### Data and Code Availability

MATLAB code for droplet analysis will be made available by G. Maryu upon its publication.

### EXPERIMENTAL MODEL AND SUBJECT DETAILS

#### Egg extract preparation

Cycling *Xenopus* egg extracts and sperm chromatin DNA were prepared as previously described [2], except that eggs were activated with calcium ionophore A23187 (200 ng/μL) (Sigma-Aldrich) rather than an electric shock [18]. The cytosolic materials were extracted via two-step centrifugation at 20,000g. Freshly prepared extracts were mixed with extract buffer, demembranated sperm chromatin, energy mix, purified proteins, and other reagents for specific experiments. In all experiments, the egg extract always accounted for 70% of the total volume, and the amount of extract buffer was adjusted when mixing multiple reagents.

### METHOD DETAILS

#### Plasmids and molecular cloning

For the cloning and expression of recombinant protein in E.coli, the pET system was employed as a backbone vector (Novagen). The cDNA of NLS-mCherry-NLS and EBFP2 were amplified by PCR and inserted into pET-45b vector plasmid with restriction enzyme. EKAREV was a gift from Dr. Michiyuki Matsuda through Addgene (Addgene, Cambridge, MA). All FRET biosensors in this study were developed by modification of this biosensor. Therefore, all FRET biosensors were composed of the same elements except for the sensor domain, ligand domain, and linker length. For replacement of sensor domain of EKAREV including FQFP ERK binding domain removed ones and negative control TA mutation was developed with linker ligation fragment that was annealed with double-stranded oligonucleotide and restriction enzymes. To utilize the same fluorophores, NLS, and affinity tag, cDNA of sensor domain and ligand domain of existing Plk1-cyclin B1-based Cdk1 sensor was amplified by PCR and introduced into the EKAREV plasmid with restriction enzymes. The coding region of the FRET biosensor in those plasmids was transferred into a pET-21a vector that has C-terminal 6x His-tag by restriction enzymes.

For mRNA purification by in vitro transcription, the pCS2+ plasmid that has an SP6 promoter region was used as a backbone vector. cDNA of Katushka and mScarlet-i were amplified by PCR and replaced with the YFP protein region of pCS2-CyclinB-YFP that we used before [18]. pmScarlet-i_C1 was a gift from Dr. Dorus Gadella through Addgene.

#### Fluorescence-Labeled Reporters

NLS-mCherry-NLS protein, FRET biosensors, and EBFP2 protein were expressed in BL21 (DE3) competent cells (New England Biolabs) that were induced by 0.4 mM IPTG (Invitrogen) overnight at 18°C. After cells were corrected by centrifugation, they were suspended by ice-cold lysis buffer (25 mM Na_2_HPO_4_, 25 mM NaH_2_PO_4_, 300 mM NaCl, 20 mM Imidazole, 1 mg/mL Lysozyme, 1 mM PMSF) and then incubated in a cold room for an hour. Cells were also broken down to release protein through sonication (Branson). Sonication condition is as follows, on ice, duty cycle: 50%, output control: 4, on:30 sec off:30 sec, and 6 cycles. His60 Ni Superflow Resin (TakaraBio) and Poly-Prep Chromatography Columns were equilibrated by a wash buffer (25 mM Na_2_HPO_4_, 25 mM NaH_2_PO_4_, 300 mM NaCl, 20 mM Imidazole) beforehand. Target proteins were incubated overnight at 4 °C and eluted with a 4 mL elution buffer (25 mM Na_2_HPO_4_, 25 mM NaH_2_PO_4_, 300 mM NaCl, 300 mM Imidazole) after three times wash by wash buffer. All purified proteins were filtered and concentrated by Amicon Ultra-4 10kDa cutoff (Millipore). The quality of purified protein was checked by CBB staining with EZBlue Gel Staining Reagent (Sigma-Aldrich), and their concentration was measured by the Bradford method with a spectrometer (DeNovix). The purified proteins were aliquoted, flash-frozen, and stored at −80°C until use.

All mRNAs were transcribed in vitro and purified using mMESSAGE mMACHINE SP6 Transcription Kit (Ambion) and RNeasy Micro Kit (QIAGEN). mRNA concentration was measured by a spectrometer.

#### Droplet formation

Fresh egg cycling extract was mixed with a surfactant on a PDMS chamber that was designed in the previous study to form droplets [20]. 2% 008-FluoroSurfactant in HFE7500 (Ran Biotechnologies, Inc.) was used as the oil phase for microfluidics. The oil and aqueous phase were driven with Elveflow OB1 multi-channel flow controller (Elveflow) and went through microbore PTFE tubings (ColeParmer) from vials to a PDMS device. For the single-channel droplet formation, the air pressure of the aqueous phase is 1.5 psi, and the oil phase is 2.0 psi. The air pressure of the aqueous phase was controlled between 0.05 psi to 1.5 psi in the reverse phase for two-channel tuning to yield droplets with different concentrations of morpholino in a wide set continuous range. This flow pump was controlled by Elveflow Software Interface. When droplets were generated and loaded into glass tubes that were pre-coated with trichloro (1H,1H,2H,2H-perfluorooctyl) silane, the tubes were immersed in a glass-bottom dish (WillCo Wells) filled with mineral oil (Macron Fine Chemicals) to prevent sample evaporations.

#### Live cell imaging

Imaging was performed with an inverted microscope IX83 (Olympus) equipped with a UPlanSApo 4x/0.16 objective lens (Olympus), an ORCA-Flash4.0 V3Digital CMOS camera (Hamamatsu Photonics), X-Cite Xylis Broad Spectrum LED Illumination System (Exelitas Technologies Corp.), and a motorized XY stage (Prior Scientific Inc.) at room temperature. The microscope was controlled by Micro-Manager software [45, 46]. Time-lapse images were recorded in bright-field and multiple fluorescence channels at a time interval of 3-6 min up to 2 days.

### QUANTIFICATION AND STATISTICAL ANALYSIS

#### Image processing

Image processing for droplet-based images was performed using custom MATLAB (MathWorks) scripts as described in a previous study [20]. Bright-field images were used for the segmentation and tracking of individual droplets. Fluorescent images of nuclear makers were used to define cellular compartments such as nuclear and cytoplasm regions. The standard deviation of fluorescent intensity, skewness of intensity histogram, and multiple thresholds were calculated in every time-point and every droplet (Figure S3A). Based on standard deviation value and skewness, a certain threshold for the Otsu method was applied for nuclear segmentation. Based on segmentation results, we calculated the average ratio value of the cytosolic and nuclear region intensities and defined the NEF, NEB Start, and NEB End points (Figure 3F, S3B-3C). Mean intensities from all fluorescent channels were calculated for each subcellular region of tracked droplets to obtain time course data. In the mitotic phase, the nuclear region and cytosolic region had the same area due to nuclear disappearance.

#### Creation of FRET/CFP ratio image stack

To generate colored FRET/CFP ratio images shown in figures, image stacks of background-subtracted CFP, FRET images were created with custom Fiji macro. After the creation of the FRET/CFP ratio image stack by image calculator function, its values were represented by the intensity-modulated display mode with custom Fiji macro (gift from Dr. Yohei Kondo).

#### Data analysis

Based on the calculated FRET/CFP ratio value and CFP intensity, peaks and troughs of Cdk1 activity were detected by custom MATLAB scripts. All peak and trough data were screened manually. In this study, from one peak to the next peak is counted as a certain oscillation. Also, the rising period and falling period are defined by peaks and troughs data. With sperm chromatin DNA condition, some droplets contain a nucleus. We defined the NEF, NEB Start, and NEB End with the cytoplasm/nuclear intensity value ratio of NLS-tagged fluorescent protein. When the cycling extract did not have a nucleus including the mitotic phase, this ratio value must be 1.0 because the nuclear and cytosolic regions had the same area. Therefore, the time point when the ratio value turned to lower than 1.0 was defined as NEF, and the time point when it returned to 1.0 again was defined as the NEB End. To determine the NEB Start, we fitted the last 10 frame data from the Cdk1 activation peak by the logistic function (Eq.1), and the 1st timepoint that estimated value is larger than minimum asymptote was defined as the NEB Start (Figure S3B and 3C).

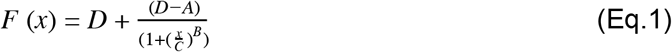

For correction of decay trends of FRET/CFP ratio values, linear baselines were estimated in each droplet with its trough values. The amplitude of Cdk1 activity was calculated by subtraction from the raw peak signal to the estimated baseline value at the time point of the peak. The average value of 1st trough values of cytoplasmic-only droplets was used for data normalization for all droplets with both raw signals and corrected signals.

To compare the Cdk1 activation shape with nucleus droplets and cytoplasmic-only droplets, all oscillation data were normalized into 250 data point vectors. A vector that had less than 250 original data points was interpolated by linearly estimated values. On the other hand, extra data points were expunged from more than 250 original data points vectors.

## Supplemental Figures

**Figure S1.**
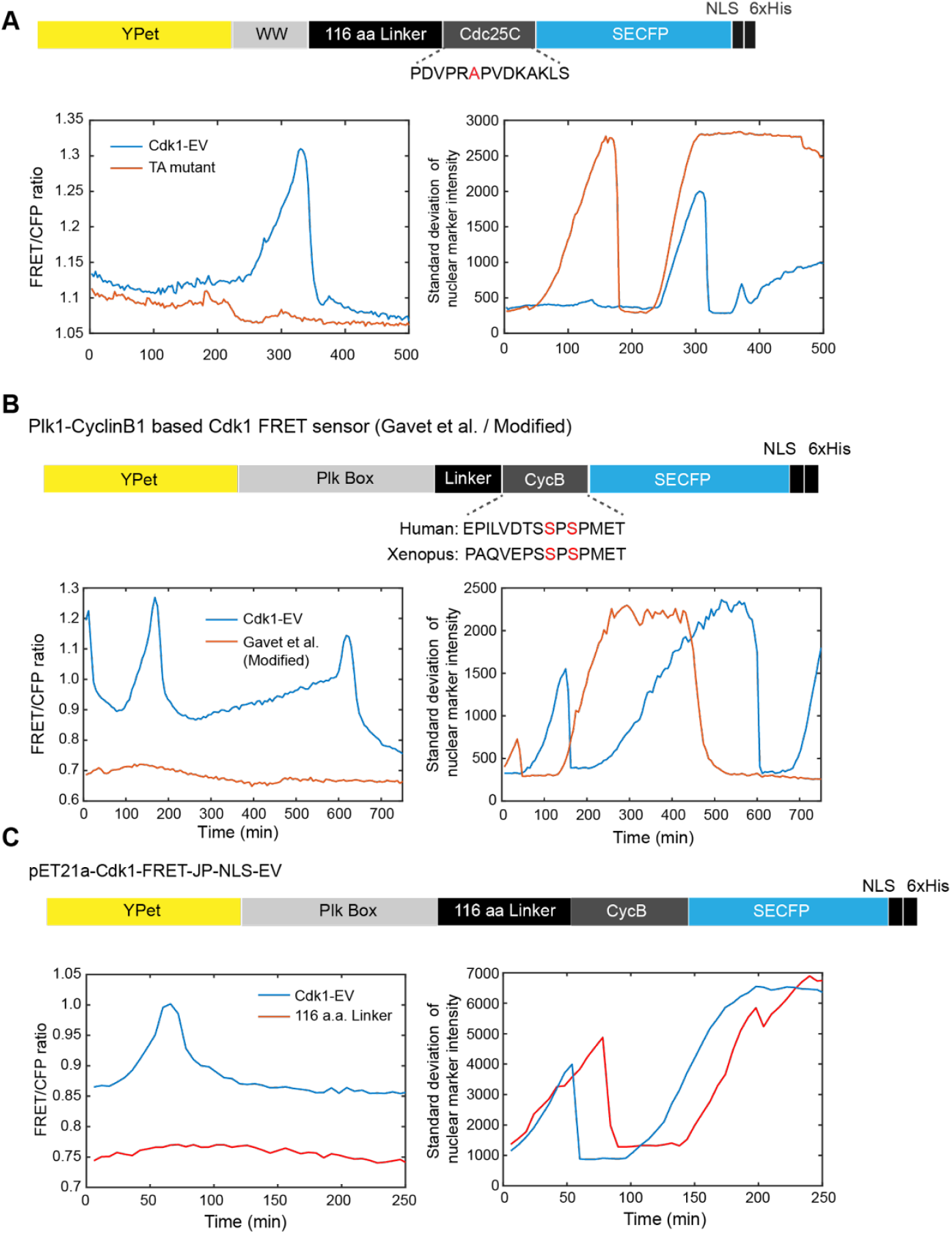
Scheme of design and representative plots for variants of the Cdk1 biosensor. (A) A comparison result of Cdk1-EV (blue line) and the phospho-dead TA mutant of Cdk1-EV (red line). Tyrosine residue in the sensor domain was replaced with Alanine as shown in the scheme. Although both representative droplets passed the mitotic phase as the standard deviation value of the nuclear marker, NLS-mCherry-NLS, showed (bottom right plot), the mutant didn’t show a FRET/CFP ratio change (bottom left plot). (B) A comparison result of response to Cdk1 activity in *Xenopus* egg extract between purified Cdk1-EV (blue line) and purified reported Cdk1 FRET biosensor (red line). The ratio value of the reported Cdk1 FRET biosensor was less changed than Cdk1-EV (bottom left). Both droplets showed a cell cycle with a nuclear marker channel (bottom right). The phosphorylation site was conserved between human and *Xenopus*, but the first half of the cyclin B1 peptide was different. This may relate to the specificity of Cdk1 activation in *Xenopus* egg extract. (C) A comparison result of Cdk1-EV (blue line) and linker length modified reported Cdk1 FRET biosensor (red line). The linker length was changed from 5 amino acid sequence to 116 amino acid sequence to improve the gain of the reported Cdk1 FRET biosensor. However, we failed to observe the Cdk1 activation with the linker modified biosensor (bottom left), although both representative droplets showed a cell cycle with a nuclear marker, NLS-mCherry-NLS (bottom right).

**Figure S2.**
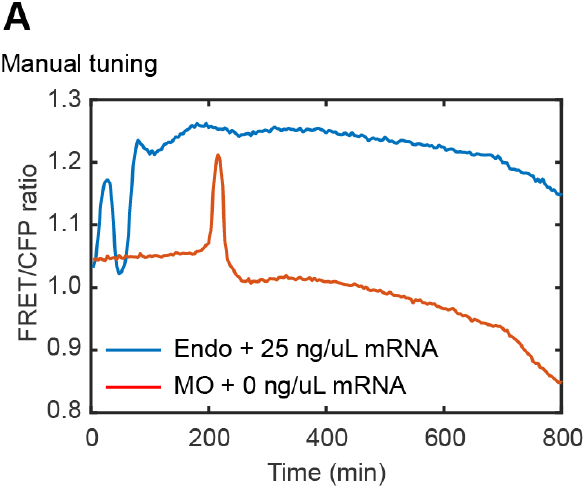
Other variations of Cdk1 activity dynamics. (A) The cell cycle oscillator of a droplet that contained endogenous mRNA and 25 ng/uL exogenous human cyclin B1 mRNA kept high Cdk1 activity after the 2nd peak (blue). On the other hand, there was a single peak and low ratio value with morpholino and no exogenous mRNA condition (red).

**Figure S3.**
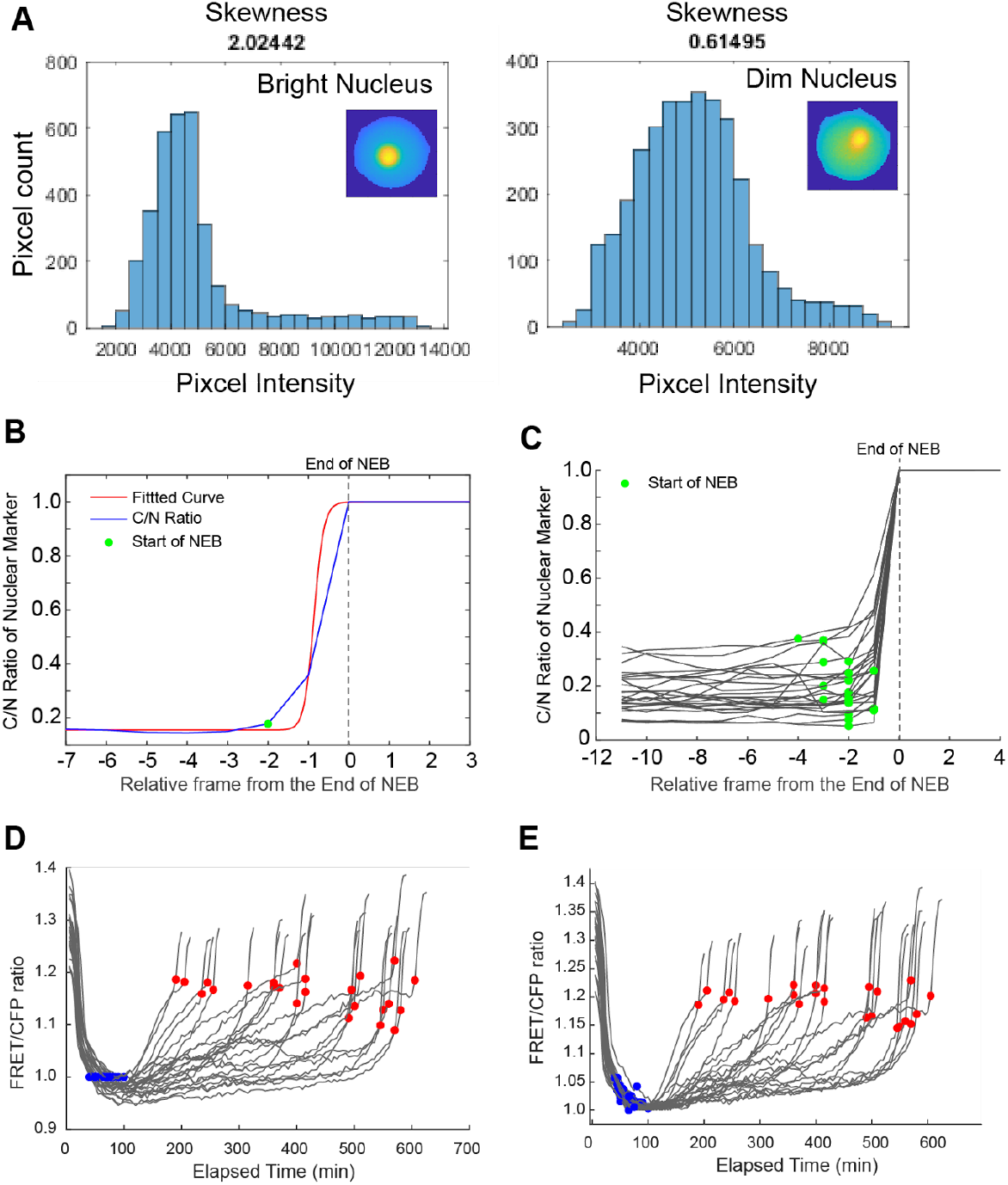
Nuclear segmentation and following analysis related nuclear envelope-related events. (A) Intensity histogram of bright nucleus droplet (left) and dim nucleus droplet (right). The color of inset droplet images indicated relative brightness in the droplets. Bright nucleus droplets showed higher skewness values with a long tail of histogram than dim ones. A different threshold value was applied based on the skewness in each droplet for nuclear segmentation. (B) Estimation of NEB Start points by logistic function fitting. The C/N ratio value of nuclear marker intensity (blue) was fitted by logistic function (red). The first C/N ratio value above fitted value was defined as NEB start as green dots. (C) Representative droplets of NEB Start result. Almost all droplets started NEB 2 or 3 frames (10~15 min) before the NEB End. This result is consistent with previous studies that took time-lapse imaging of NEB with human culture cells [S1, S2]. (D) Cdk1 activities shown in Figure 3H were normalized with the ratio value at NEF. The ratio values at the NEB end were similar among different period lengths. (E) Cdk1 activities shown in Figure 3H were normalized with the ratio value at the trough of its cycle. The ratio values at the NEB end were similar among different period lengths.

**Figure S4.**
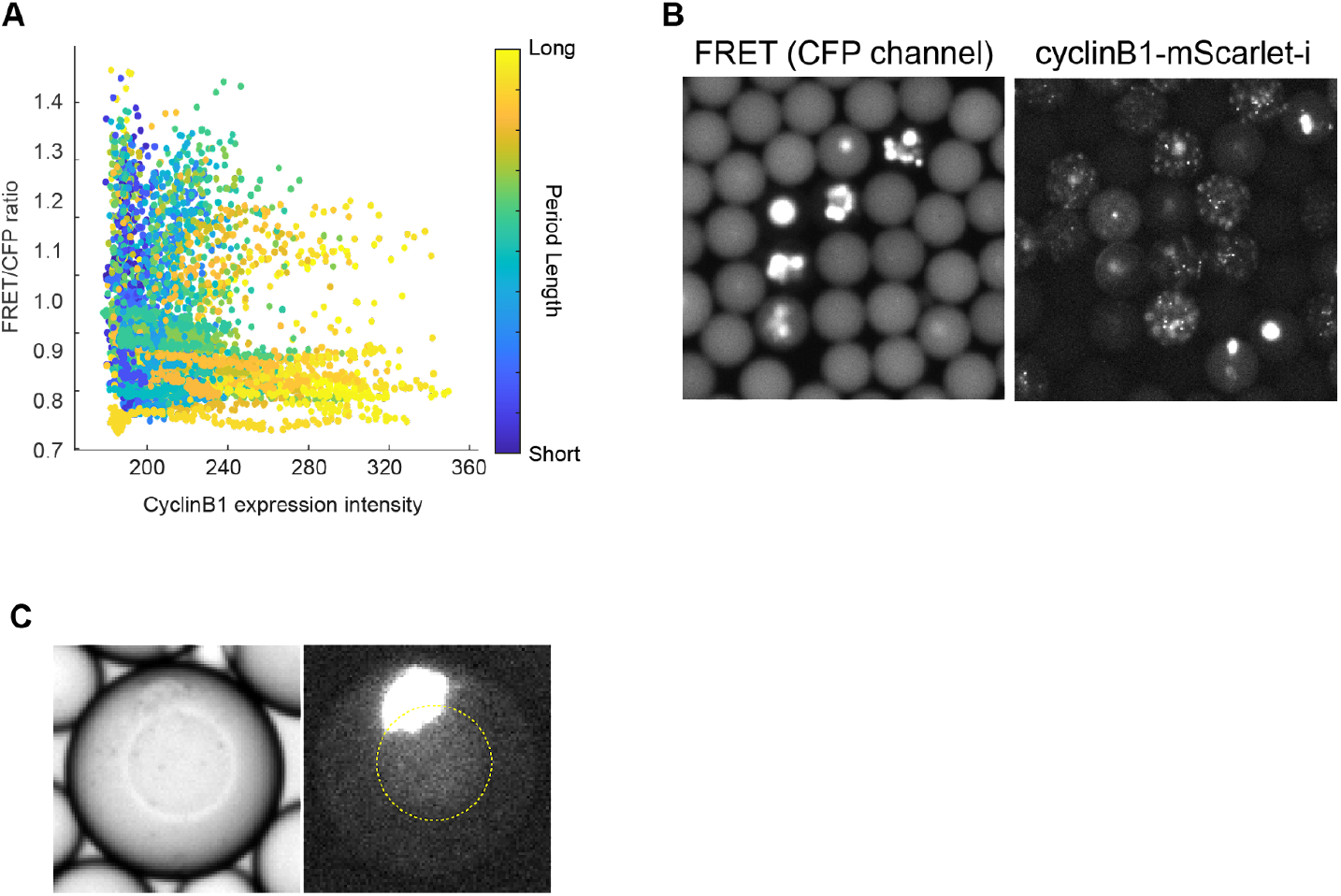
Phase-plane trajectories and cyclin B1 puncta. (A) Phase-plane trajectories of cyclin B1 expression and raw FRET/CFP ratio value. The color bar indicates period length. This plot contains all cycles that were observed in different droplets, different cycles, and both nuclear conditions. (B) Puncta structure is specifically observed in cyclin B1 channel, not CFP channel for FRET biosensor. (C) Representative droplet which had a bright large structure localized right beside the nucleus. A sphere at the center of the bright field image (left) is a nucleus. A bright structure and dim nucleus-like shape localization were observed in the fluorescent channel (right). A nucleus position was indicated by a yellow circle.

## Supplemental Video

**Video S1. Visualization of Cdk1 oscillation in cell-sized microemulsion droplets without nuclei.** Droplets that contain *Xenopus* egg extract and purified Cdk1-EV protein were formed in a microfluidic device. The color in droplets indicates Cdk1 activity. The warm color indicates high activity and the cold color is for low activity. The ratio range of this video was 1.25-0.8.

**Video S2. Visualization of Cdk1 activity and nuclear maker in droplets with nuclei.**

Sperm chromatin DNA was added to form nuclei in the droplets. Nuclear droplets and cytoplasmic-only droplets were observed in the same field of view. By the benefit of NLS, Cdk1-EV localized in nuclei and enabled us to measure nuclear Cdk1 activity. Cdk1 was activated not only in the mitotic phase but also interphase. The ratio range of this video was 1.1-0.65.

**Video S3. Simultaneous imaging of Cdk1-EV and cyclin B1-mScarlet-i expression.**

Expression and degradation of cyclin B1 was visualized with Cdk1-EV. This video corresponds to Figures 4A–4C. The ratio range of this video was 0.9-0.7.

**Video S4. Comparison of cyclin B1 degradation processes between droplets with or without puncta.**

This video corresponds to Figure 4H–4J. The time frames of these droplets were aligned at the NEB point (00:00 in the video).

## References

1. Hahn, H. S., Nitzan, A., Ortoleva, P., and Ross, J. (1974). Threshold excitations, relaxation oscillations, and effect of noise in an enzyme reaction. Proc. Natl. Acad. Sci. USA 71, 4067–4071.

2. Murray, A. W. (1991). Cell cycle extracts. Methods Cell Biol. 36, 581–605.

3. Sha, W., Moore, J., Chen, K., Lassaletta, A. D., Yi, C.-S., Tyson, J. J., and Sible, J. C. (2003). Hysteresis drives cell-cycle transitions in Xenopus laevis egg extracts. Proc. Natl. Acad. Sci. USA 100, 975–980.

4. Tsai, T. Y.-C., Choi, Y. S., Ma, W., Pomerening, J. R., Tang, C., and Ferrell, J. E. (2008). Robust, tunable biological oscillations from interlinked positive and negative feedback loops. Science 321, 126–129.

5. Yang, Q., and Ferrell, J. E. (2013). The Cdk1-APC/C cell cycle oscillator circuit functions as a time-delayed, ultrasensitive switch. Nat. Cell Biol. 15, 519–525.

6. Li, Z., Wang, S., Sun, M., Jin, M., Khain, D., and Yang, Q. (2021). High-resolution mapping of the period landscape reveals polymorphism in cell cycle frequency tuning. BioRxiv.

7. Santos, S. D. M., Wollman, R., Meyer, T., and Ferrell, J. E. (2012). Spatial positive feedback at the onset of mitosis. Cell 149, 1500–1513.

8. Gavet, O., and Pines, J. (2010). Activation of cyclin B1-Cdk1 synchronizes events in the nucleus and the cytoplasm at mitosis. J. Cell Biol. 189, 247–259.

9. Gavet, O., and Pines, J. (2010). Progressive activation of CyclinB1-Cdk1 coordinates entry to mitosis. Dev. Cell 18, 533–543.

10. Levasseur, M. D., Thomas, C., Davies, O. R., Higgins, J. M. G., and Madgwick, S. (2019). Aneuploidy in Oocytes Is Prevented by Sustained CDK1 Activity through Degron Masking in Cyclin B1. Dev. Cell 48, 672–684.e5.

11. Deneke, V. E., Melbinger, A., Vergassola, M., and Di Talia, S. (2016). Waves of cdk1 activity in S phase synchronize the cell cycle in drosophila embryos. Dev. Cell 38, 399–412.

12. Belal, A. S. F., Sell, B. R., Hoi, H., Davidson, M. W., and Campbell, R. E. (2014). Optimization of a genetically encoded biosensor for cyclin B1-cyclin dependent kinase 1. Mol. Biosyst. 10, 191–195.

13. Komatsu, N., Aoki, K., Yamada, M., Yukinaga, H., Fujita, Y., Kamioka, Y., and Matsuda, M. (2011). Development of an optimized backbone of FRET biosensors for kinases and GTPases. Mol. Biol. Cell 22, 4647–4656.

14. Harvey, C. D., Ehrhardt, A. G., Cellurale, C., Zhong, H., Yasuda, R., Davis, R. J., and Svoboda, K. (2008). A genetically encoded fluorescent sensor of ERK activity. Proc. Natl. Acad. Sci. USA 105, 19264–19269.

15. Franckhauser, C., Mamaeva, D., Heron-Milhavet, L., Fernandez, A., and Lamb, N. J. C. (2010). Distinct pools of cdc25C are phosphorylated on specific TP sites and differentially localized in human mitotic cells. PLoS One 5, e11798.

16. Bonnet, J., Mayonove, P., and Morris, M. C. (2008). Differential phosphorylation of Cdc25C phosphatase in mitosis. Biochem. Biophys. Res. Commun. 370, 483–488.

17. Aoki, K., Kumagai, Y., Sakurai, A., Komatsu, N., Fujita, Y., Shionyu, C., and Matsuda, M. (2013). Stochastic ERK activation induced by noise and cell-to-cell propagation regulates cell density-dependent proliferation. Mol. Cell 52, 529–540.

18. Guan, Y., Li, Z., Wang, S., Barnes, P. M., Liu, X., Xu, H., Jin, M., Liu, A. P., and Yang, Q. (2018). A robust and tunable mitotic oscillator in artificial cells. Elife 7.

19. Guan, Y., Wang, S., Jin, M., Xu, H., and Yang, Q. (2018). Reconstitution of Cell-cycle Oscillations in Microemulsions of Cell-free Xenopus Egg Extracts. J. Vis. Exp.

20. Sun, M., Li, Z., Wang, S., Maryu, G., and Yang, Q. (2019). Building Dynamic Cellular Machineries in Droplet-Based Artificial Cells with Single-Droplet Tracking and Analysis. Anal. Chem. 91, 9813–9818.

21. Zhang, W., and Liu, H. T. (2002). MAPK signal pathways in the regulation of cell proliferation in mammalian cells. Cell Res. 12, 9–18.

22. Albeck, J. G., Mills, G. B., and Brugge, J. S. (2013). Frequency-modulated pulses of ERK activity transmit quantitative proliferation signals. Mol. Cell 49, 249–261.

23. Trunnell, N. B., Poon, A. C., Kim, S. Y., and Ferrell, J. E. (2011). Ultrasensitivity in the regulation of cdc25c by cdk1. Mol. Cell 41, 263–274.

24. Kim, S. Y., and Ferrell, J. E. (2007). Substrate competition as a source of ultrasensitivity in the inactivation of Wee1. Cell 128, 1133–1145.

25. Han, Z., Yang, L., MacLellan, W. R., Weiss, J. N., and Qu, Z. (2005). Hysteresis and cell cycle transitions: how crucial is it? Biophys. J. 88, 1626–1634.

26. Weitz, M., Kim, J., Kapsner, K., Winfree, E., Franco, E., and Simmel, F. C. (2014). Diversity in the dynamical behaviour of a compartmentalized programmable biochemical oscillator. Nat. Chem. 6, 295–302.

27. Cao, Y., Wang, H., Ouyang, Q., and Tu, Y. (2015). The free energy cost of accurate biochemical oscillations. Nat. Phys. 11, 772–778.

28. Son, S., Kang, J. H., Oh, S., Kirschner, M. W., Mitchison, T. J., and Manalis, S. (2015). Resonant microchannel volume and mass measurements show that suspended cells swell during mitosis. J. Cell Biol. 211, 757–763.

29. Zlotek-Zlotkiewicz, E., Monnier, S., Cappello, G., Le Berre, M., and Piel, M. (2015). Optical volume and mass measurements show that mammalian cells swell during mitosis. J. Cell Biol. 211, 765–774.

30. Novak, B., Csikasz-Nagy, A., Gyorffy, B., Chen, K., and Tyson, J. J. (1998). Mathematical model of the fission yeast cell cycle with checkpoint controls at the G1/S, G2/M and metaphase/anaphase transitions. Biophys Chem 72, 185–200.

31. Stern, B., and Nurse, P. (1996). A quantitative model for the cdc2 control of S phase and mitosis in fission yeast. Trends Genet. 12, 345–350.

32. Lindqvist, A., van Zon, W., Karlsson Rosenthal, C., and Wolthuis, R. M. F. (2007). Cyclin B1-Cdk1 activation continues after centrosome separation to control mitotic progression. PLoS Biol. 5, e123.

33. Lindqvist, A. (2010). Cyclin B-Cdk1 activates its own pump to get into the nucleus. J. Cell Biol. 189, 197–199.

34. Falahati, H., Hur, W., Di Talia, S., and Wieschaus, E. (2021). Temperature-Induced uncoupling of cell cycle regulators. Dev. Biol. 470, 147–153.

35. Swaffer, M. P., Jones, A. W., Flynn, H. R., Snijders, A. P., and Nurse, P. (2016). CDK substrate phosphorylation and ordering the cell cycle. Cell 167, 1750–1761.e16.

36. Basu, S., Roberts, E. L., Jones, A. W., Swaffer, M. P., Snijders, A. P., and Nurse, P. (2020). The hydrophobic patch directs cyclin B to centrosomes to promote global CDK phosphorylation at mitosis. Curr. Biol. 30, 883–892.e4.

37. Moore, J. D., Kirk, J. A., and Hunt, T. (2003). Unmasking the S-phase-promoting potential of cyclin B1. Science 300, 987–990.

38. Pomerening, J. R., Kim, S. Y., and Ferrell, J. E. (2005). Systems-level dissection of the cell-cycle oscillator: bypassing positive feedback produces damped oscillations. Cell 122, 565–578.

39. Chang, D. C., Xu, N., and Luo, K. Q. (2003). Degradation of cyclin B is required for the onset of anaphase in Mammalian cells. J. Biol. Chem. 278, 37865–37873.

40. Terasaki, M., Okumura, E.-I., Hinkle, B., and Kishimoto, T. (2003). Localization and dynamics of Cdc2-cyclin B during meiotic reinitiation in starfish oocytes. Mol. Biol. Cell 14, 4685–4694.

41. Chen, Q., Zhang, X., Jiang, Q., Clarke, P. R., and Zhang, C. (2008). Cyclin B1 is localized to unattached kinetochores and contributes to efficient microtubule attachment and proper chromosome alignment during mitosis. Cell Res. 18, 268–280.

42. Bentley, A. M., Normand, G., Hoyt, J., and King, R. W. (2007). Distinct sequence elements of cyclin B1 promote localization to chromatin, centrosomes, and kinetochores during mitosis. Mol. Biol. Cell 18, 4847–4858.

43. Araujo, A. R., Gelens, L., Sheriff, R. S. M., and Santos, S. D. M. (2016). Positive Feedback Keeps Duration of Mitosis Temporally Insulated from Upstream Cell-Cycle Events. Mol. Cell 64, 362–375.

44. Schindelin, J., Arganda-Carreras, I., Frise, E., Kaynig, V., Longair, M., Pietzsch, T., Preibisch, S., Rueden, C., Saalfeld, S., Schmid, B., et al. (2012). Fiji: an open-source platform for biological-image analysis. Nat. Methods 9, 676–682.

45. Edelstein, A. D., Tsuchida, M. A., Amodaj, N., Pinkard, H., Vale, R. D., and Stuurman, N. (2014). Advanced methods of microscope control using μManager software. J. Biol. Methods 1.

46. Edelstein, A., Amodaj, N., Hoover, K., Vale, R., and Stuurman, N. (2010). Computer control of microscopes using μManager. Curr. Protoc. Mol. Biol. Chapter 14, Unit14.20.

## Supplemental reference

S1. Beaudouin, J., Gerlich, D., Daigle, N., Eils, R., and Ellenberg, J. (2002). Nuclear envelope breakdown proceeds by microtubule-induced tearing of the lamina. Cell 108, 83–96.

S2. Shankaran, S. S., Mackay, D. R., and Ullman, K. S. (2013). A time-lapse imaging assay to study nuclear envelope breakdown. Methods Mol. Biol. 931, 111–122.

